# Adjunctive nebulization of Idursulfase to intravenous treatment enhances cardiac proteome adaptations in IDS-KO mice

**DOI:** 10.64898/2026.09.07.749889

**Authors:** Vinicius M. Mariani, Anatália Labilloy, W. Todd Cade, Paulo R. Ferreira, Leonardo F. Ferreira

## Abstract

Mucopolysaccharidosis Type II (MPS-II) is a rare X-linked lysosomal storage disorder, characterized by deficiency of the lysosomal hydrolase iduronate 2-sulphatase (IDS). Cardiac impairments contribute to mortality in patients. Enzyme replacement therapy (ERT) with Idursulfase is a promising treatment for the cardiac dysfunction. We compared the proteome changes following intravenous (IV) with and without adjunct nebulized ERT in IDS-KO mice. Male mice were assigned to 4 groups: 1) WT IV + nebulized saline (WT; n=8); 2) IDS-KO IV + nebulized saline (KO; n=5); 3) IDS-KO IV Idursulfase (1 mg/kg; KO-IV, n=5); 4) IDS-KO IV (1 mg/kg) + nebulized Idursulfase (0.33 mg/mouse; KO-NEB, n=5). Treatments were administered once a week starting at 8 weeks of age and the heart was harvested 12 weeks post-ERT. Mouse heart samples were prepared for global proteomics measurements with Nano-liquid chromatography tandem mass spectrometry (n=3/group; Nano-LC/MS/MS). Global limma moderated F-test analysis identified 928 differently abundant proteins across the four groups. Pairwise Euclidean distances from the IDS-KO were significantly greater for the KO- NEB vs KO-IV group (47.5 ± 3.7 vs 40.8 ± 1.1; p < 0.001), showing that adjunct nebulization elicited a more profound proteomic shift than IV alone. The mean z-score compartment analysis revealed that the subcellular compartment protein profile of KO-NEB shifted towards the WT profile, whereas this pattern was less evident in KO-IV. The cluster analysis also revealed treatment-specific and rescue-like protein modules. Overall, our results suggest that adjunctive nebulized Idursulfase ERT enhances the cardiac proteomic changes caused by IV treatment in IDS-KO mice. Proteomics data are available via ProteomeXchange with identifier PXD083242.

**Synopsis:** Adjunctive nebulized Idursulfase ERT enhances the cardiac proteomic changes caused by IV treatment in IDS-KO mice.

## 1 Introduction

Mucopolysaccharidosis Type II (MPS-II), also known as Hunter syndrome, is a rare X- linked genetic disorder with an incidence rate of 0.38 to 1.09 per 100,000 live male births [1]. The disorder is characterized by a deficiency of the lysosomal hydrolase iduronate 2- sulphatase (IDS) enzyme, which is critical for the breakdown of glycosaminoglycans (GAGs) heparan sulfate and dermatan sulfate [1]. IDS deficiency and consequent accumulation of GAGs trigger a series of multisystemic impairments that affect the liver, spleen, bones, joints, respiratory tract, and heart [2].

Cardiac dysfunction is a major driver of mortality in MPS-II patients. MPS-II cardiomyopathy involves valve insufficiency and remodeling [2, 3], electrocardiogram alterations [4, 5], and left ventricular hypertrophy and dysfunction [3, 6, 7]. These clinical manifestations can culminate in severe heart failure and premature death [8]. Data from the pre-clinical model of MPS-II, IDS knockout (KO) mice, support impaired cardiac mitochondria respiration, reduced activity of key enzymes involved in metabolism and disrupted mitochondria-lysosome interaction as contributing mechanisms to the cardiac phenotype in MPS-II [9]. However, it is unclear how MPS-II reshapes the cardiac proteome. Proteomics profiling of the heart can lend insights into mechanisms of cardiomyopathy and identify targets for preventing and reversing cardiac impairments.

Intravenous delivery of the purified recombinant form of IDS, Idursulfase (ELAPRASE®, Takeda, Lexington, MA) is an FDA-approved treatment that delays the onset of cardiac valve disease, decreases left ventricle mass index, and increases forced vital capacity and six- minute walk distance [10–12]. Despite these reported benefits of ERT, cardiorespiratory failure is still the predominant cause of death in treated and untreated patients [13–15]. Results from pre-clinical rodent studies indicate that intravenous Idursulfase absorption is heterogeneous among organs, with the liver absorbing 30-40% of the delivered dose, which equates to a 60- fold higher uptake compared to the heart [16]. Therefore, approaches to enhance ERT delivery to the heart are needed in MPS-II. Nebulized Idursulfase ERT treatment is an alternative route for enhancing delivery to the cardiorespiratory system in IDS-KO mice [17]. Nebulized drug administration elicits a local increase in drug concentration in the lungs that is rapidly transferred to the heart and subsequently to the circulation, bypassing the liver filtering system [18]. However, the effects of IV and nebulized Idursulfase treatment on the heart of IDS-KO mouse are unclear. The goal of our study was to define the cardiac proteome features of IDS deficiency and changes elicited by IV Idursulfase ERT with adjunct nebulization delivery.

## 2 |Methodology

### 2.1 Animals and Procedures

All animal procedures were described in detail in our previous study [17]. Male wildtype (WT) C57BL/6N mice and IDS-knockout (KO) mice (024744, B6N.Cg-Idstm1Muen/J; Jackson Laboratory, Bar Harbor, ME) were housed on a 12 h:12 h light-dark cycle with access to chow and water ad libitum. We studied only male mice since MPS-II is a X-linked disease affecting almost exclusively males.

Mice were assigned to one of the four following groups: 1) WT IV + nebulized saline (WT); 2) IDS-KO IV + nebulized saline (KO); 3) IDS-KO IV Idursulfase (1 mg/kg; KO-IV); 4) IDS-KO IV (1 mg/kg) + adjunctive nebulized Idursulfase (0.33 mg/mouse; KO-NEB). Treatments were administered once a week starting at 8 weeks of age. Intravenous ERT was delivered via tail vein injection while idursulfase was nebulized as 167 µL of a 2 mg/ml solution. Terminal procedures occurred 12 weeks after treatment onset and 7 days after the final dose. Mice were euthanized under anesthesia with isoflurane (5% induction, 2-3% maintenance), and a laparotomy and thoracotomy were performed to collect the hearts. We prepared samples from n = 3/group for global proteomics.

### 2.2 Cardiac Sample Preparation

Immediately after tissue harvesting, heart samples were rinsed in PBS, blotted dry, flash-frozen in liquid nitrogen (N₂), and stored at −80 °C until processing. Heart global proteomics analysis with Nano-liquid chromatography tandem mass spectrometry (Nano- LC/MS/MS) was conducted by the University of Florida Proteomics and Mass Spectroscopy Core.

Protein digestion and extraction were performed with EasyPep^TM^ MS Sample Prep Kit (Thermo Fisher Scientific). We used Qubit analysis to determine the sample volume required to yield 100 µg of final total protein. The samples were then digested with a sequencing grade trypsin/lys C rapid digestion kit (Progema) following the manufacturer’s protocol. Final digestion buffer and sample volume ratio was 3:1. Samples were incubated at 56 °C with 1 µL of 0.1 M DTT in 100 mM ammonium bicarbonate for 30 minutes, followed by the addition of 0.54 µL of 55 mM Iodoacetamide in 100 mM ammonium bicarbonate and incubation for 30 minutes at room temperature in the dark. Trypsin/lys C (1µg/µl concentration) was added to the samples and incubated at 70 °C for 1 hour. Digestion was interrupted by the addition of 0.5% TFA.

### 2.3 Global Label-free Proteomic

A Q Exactive HF Orbitrap mass spectrometer (Thermo Fisher Scientific) was used to perform Nano-LC/MS/MS. The system was operated in positive ion mode and with an UltiMate^TM^ 3000 RSLCano system. Mobile phase A was 0.1% formic acid in H_2_O and phase B (B) was acetonitrile. The phase A used for the loading pump was 0.1% trifluoroacetic acid in H_2_O. Processed samples were injected into a Pharma Fluidics µPAC^TM^ C18 trapping column (flow rate of 10 µL/mL for 3 min), then washed with 1% phase B. Chromatographic separation was performed at 40 °C. The flow rate was sustained at 750 nL/min for 15 minutes and moved afterward to 300 nL/min. Peptides were eluted off the column into the Q Exactive HF Orbitrap system with a 1% to 20% phase B gradient for 100 minutes, followed by 45% phase B for 20 minutes.

The scan sequence used to run the mass spectrometer was based on the original TopTen^TM^ method with the full scan ranging from 375 to 1575 Da at 60,000 and MS/MS scan at 15,000 resolution. Amino acid sequence was determined in consecutive scans of the 15 most abundant peaks in the spectrum. Full scan included 300,000 AGC target ion number and 50 ms injection time, and MS/MS mode included 20,000 AGC target ion number with 55 ms injection time. (N)CE/stepped NCE was set to 28. Single charged ions were excluded from MS/MS analysis. Siloxane background peak at 445.12003 was used to correct real-time mass shifts and improve mass and peptide identification accuracy.

The MS/MS spectra were analyzed with Sequest (Thermo Fisher Scientific; version IseNode in Proteome Discoverer 3.0.1.27). Searches with Sequest were performed against the mouse database with trypsin specificity. Mass tolerances were set to 10.0 ppm for precursor ions and 0.020 Da for fragment ions.

### 2.4 Data Analysis and Statistics

Normalized protein abundance values were obtained from the Proteome Discoverer output and extracted from 12 samples distributed across the 4 experimental groups: WT (F10- F12), untreated IDS-KO (KO; F1-F3), IDS-KO treated with IV IDS (KO-IV; F4-F6), and IDS-KO treated with IV and NEB IDS (KO-NEB; F7-F9). Only proteins that were detected in all three biological replicates from at least one experimental group (100% feature) were considered for analysis to enhance confidence in outcomes with n = 3/group. Proteins were also removed from the analysis when annotated in the Contaminant column of the Proteome Discoverer output.

We implemented a low-intensity, left-censored imputation strategy, applied independently to each sample to account for low abundance. We continued with linear models for microarray (limma) analysis because this approach relies on variance across all proteins, reducing false-positive results in studies with small sample sizes [19, 20]. Missing values were imputed using a Perseus-style left-censored strategy. Values were sampled from a normal distribution centered 1.8 standard deviations below the observed mean, with a standard deviation equal to 0.3 times the observed sample standard deviation. These parameters correspond to the default Perseus implementation and are used to model proteins likely present below the detection limit while minimizing distortion of the observed intensity distribution [21, 22]. To ensure reproducibility, a fixed random seed was used for imputation. Then, the log_2_-transformed and imputed abundance matrix served as input for the limma analysis. Each experimental group was encoded as a four-level categorical factor. A one-factor linear model with no intercept was fitted using the design matrix ∼0 + group, allowing one main abundance coefficient to be estimated for each experimental group.

We performed a group-effect analysis to identify alterations in protein abundance across the experimental groups. We tested three contrasts against KO: WT – KO, KO-IV – KO, and KO-NEB – KO. The resulting limma moderated F-test value defined whether any of the differences among groups were non-zero for each protein. The Benjamini-Hochberg false discovery rate (FDR) test was used to adjust the nominal p-values for all tested proteins. Proteins with FDR <0.05—significant group-associated abundance variation—were considered for downstream dimensionality reduction, structure, clustering, and annotation analyses. The planned pairwise contrasts were then evaluated using the same one-factor model: WT – KO, KO-IV – KO, and KO-NEB – KO. Therefore, positive and negative log_2_ fold-changes imply higher and lower protein abundance, respectively, in WT, KO-IV, or KO-NEB, relative to KO. To stabilize protein-wise variance estimates, we applied an empirical Bayes moderation test after contrast fitting. Again, for each contrast, p-values were adjusted across all tested proteins using the Benjamini-Hochberg FDR. We considered protein abundances statistically different in pairwise comparisons when they met the following criteria: an absolute log_2_ fold- change ≥ 1 and an adjusted p-value < 0.05.

We calculated pairwise Euclidean distances to estimate sample variance within and between groups. Samples were represented as observations and proteins as features. Protein feature was standardized across the 12 samples to have a ‘0’ mean and unit variance before distance calculation to ensure that distances reflected relative abundance changes across samples instead of differences in absolute protein intensity.

The resulting distance matrix was used both to quantify within- and between-group variation and as input for metric multidimensional scaling (MDS), implemented using the SMACOF algorithm, allowing visualization of sample relationships while preserving pairwise distances in a two-dimensional space as close as possible. To compare the magnitude of proteomic shifts between treatments, pairwise Euclidean distances from the untreated KO baseline to each treatment group were extracted and compared using a two-sided Mann- Whitney U test.

To define coordinated protein abundance changes among groups, all proteins with significant group-associated abundance changes were clustered based on their k-mean. The global protein clusters were intersected with the limma contrast results to determine how many clusters were represented in each pairwise comparison and which clusters were most represented. For downstream biological interpretation, we identified k = 6 relevant clusters (C1-C6). The final k-means model was fitted using k = 6, 100 random initializations, and a fixed random seed. The generated input matrix consists of protein-level z-scores derived from the

log_2_ –transformed and imputed abundance matrix. The clustering analysis was based on relative abundance changes across samples rather than on absolute protein intensity. For each cluster, mean protein-level z-score profiles were calculated across individual samples and experimental groups. Cluster labels were assigned to each protein and merged with protein annotation and global limma F-test statistics.

We also analyzed the localization of altered proteins to identify cellular targets of the disease and treatments. Protein accessions were queried against UniProtKB via the UniProt REST API, restricting the search to Mus musculus proteins with organism identifier 10090. To summarize localization outcomes, UnitProt subcellular location comments, Gene Ontology (GO) Cellular Component annotations, and GO-identified fields were merged into a single text for each protein. We then used keyword-based searches to identify proteins associated with selected localization categories.

The mass spectrometry proteomics data have been deposited to the ProteomeXchange Consortium via the PRIDE [23] partner repository with the dataset identifier PXD083242 and 10.6019/PXD083242.

## 3 Results

### 3.1 Protein Abundance and Cardiac Proteome Changes

Our analysis revealed a total of 2,318 proteins. Of the 2,318 proteins, 928 were identified as significant in the global limma moderated F-test. The global proteomic signature change across groups and individual samples is illustrated in **Figure 1**. We observed a total of 1,150 significant contrast-level protein abundance changes, corresponding to 757 unique proteins identified in at least one contrast: WT relative to IDS-KO, 438 proteins (up = 104, down = 334); KO-IV relative to IDS-KO, 308 proteins (up = 141, down = 167); and KO-NEB relative to IDS-KO, 404 proteins (up = 144, down = 260). The Venn diagram generated from the significant limma contrast protein sets showed that IV and NEB treatments share some similar protein, but the NEB treatment was associated with 165 unique protein changes versus 85 proteins in the IV treatment (**Figure 2**). Moreover, the KO-NEB vs IDS-KO contrast showed overlap of 75 proteins with the WT vs IDS-KO contrast vs 59 proteins for KO-IV vs IDS-KO.

**Figure 1.**
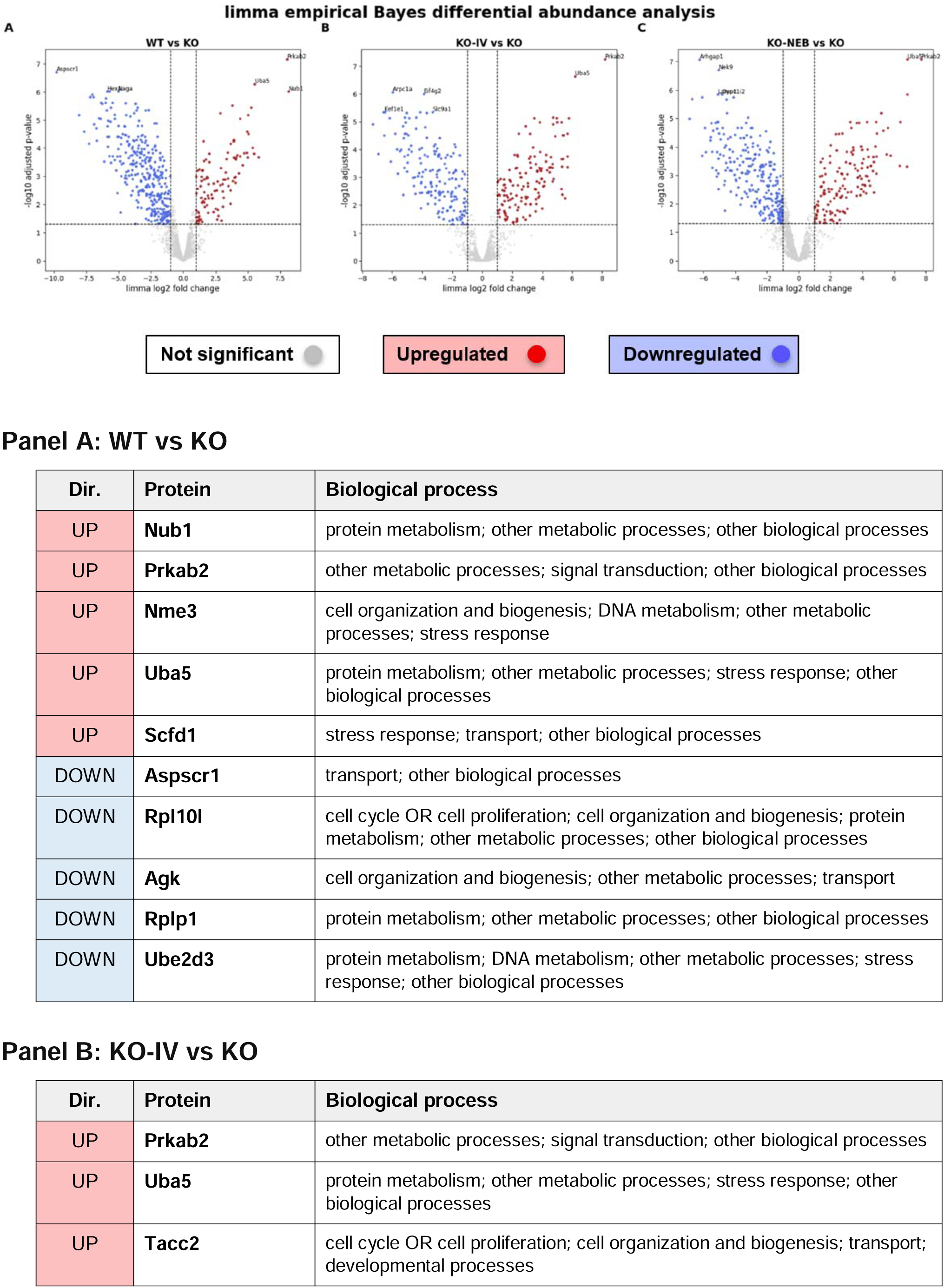
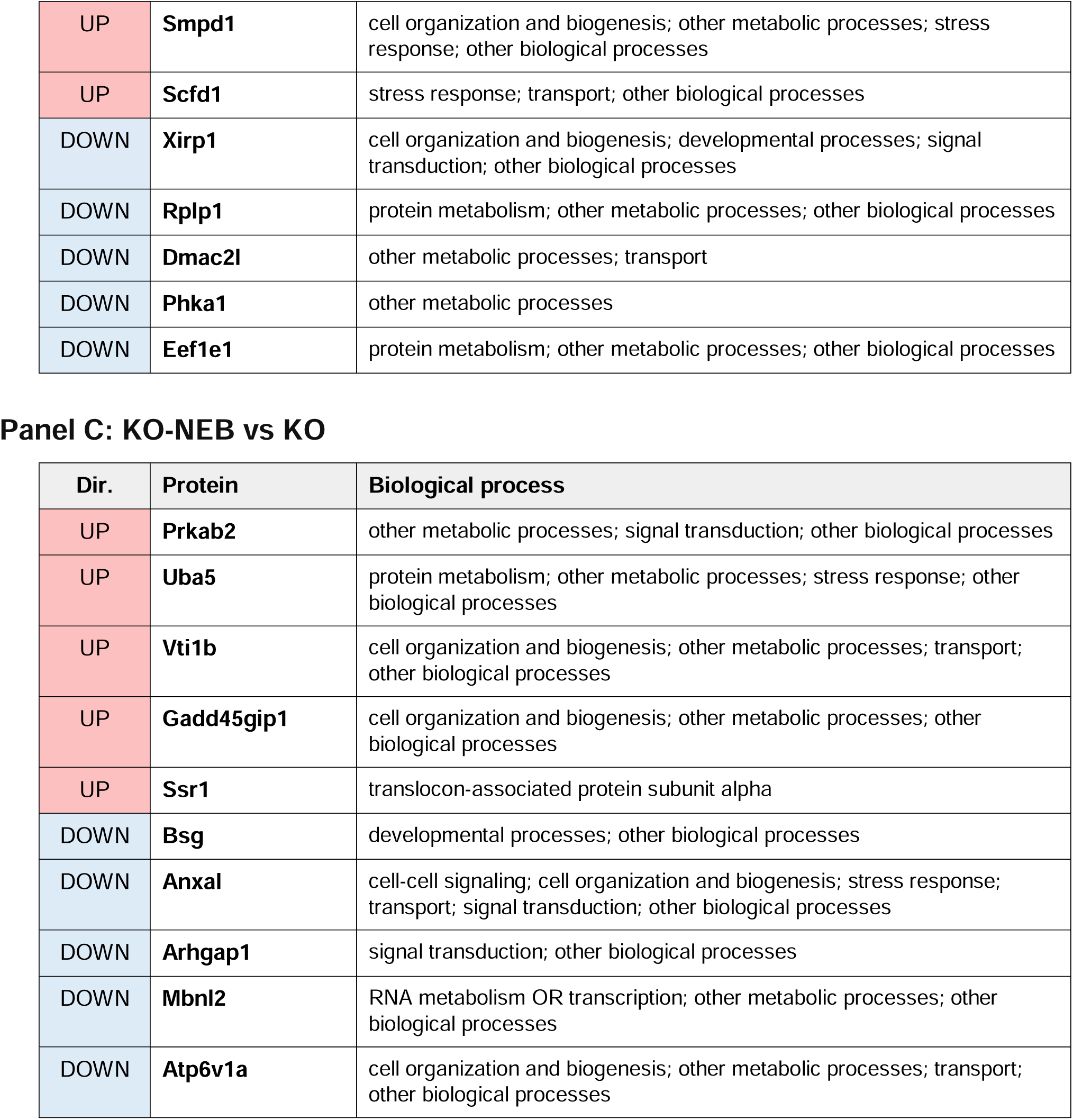
Limma empirical Bayes differential abundance analysis of cardiac proteins across pairwise contrasts. Volcano plots show differential protein abundance for WT vs KO (A), KO-IV vs KO (B), and KO-NEB vs KO (C). Each point represents one protein, with log2 fold change shown on the x-axis and −log10 adjusted p-value on the y-axis. Positive log2 fold-change values indicate higher abundance in WT, KO-IV, or KO-NEB relative to untreated IDS-KO, whereas negative values indicate lower abundance relative to IDS-KO. Dashed lines indicate the significance thresholds of |log2FC| ≥ 1 and adjusted p-value < 0.05. Significantly upregulated and downregulated proteins are shown in red and blue, respectively, while non- significant proteins are shown in gray. Protein labels indicate selected significant proteins with the strongest statistical evidence. Tables summarize the top 5 upregulated and top 5 downregulated proteins for each contrast, ranked by log2 fold-change magnitude, with biological process annotations retained from the source table.

**Figure 2.**
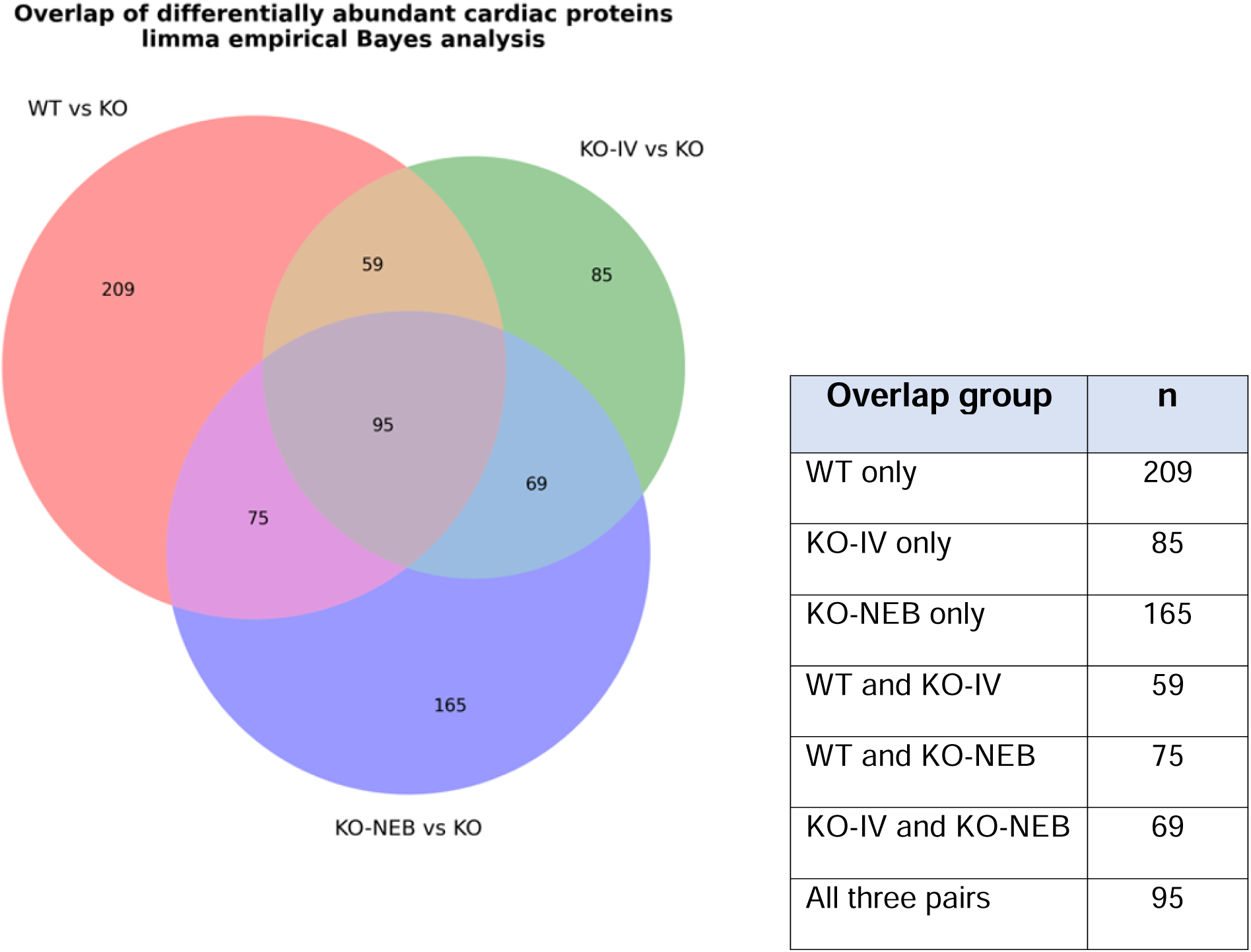
Overlap of differentially abundant cardiac proteins across pairwise contrasts. Venn diagram showing the overlap among proteins identified as differentially abundant in the planned limma contrasts WT vs KO, KO-IV vs KO, and KO-NEB vs KO. Differential abundance was defined using |log2FC| ≥ 1 and adjusted p-value < 0.05. The numbers indicate proteins unique to each contrast or shared across contrasts. KO-NEB vs KO showed a larger unique differential protein signature than KO-IV vs KO, and also showed greater overlap with WT vs KO than did KO-IV vs KO. The table summarizes the number of proteins in each overlap category.

The top 5 up- and down-regulated proteins from each pairwise contrast indicate candidate proteins associated with the disease and treatment-related proteomic signatures (**Figure 1**). For the WT vs IDS-KO comparison, Nub1, Prkab2, Nme3, Uba5 and Scfd1 were upregulated, and Aspscr1, Rpl10l, Agk, Rplp1, and Ube2d3 were downregulated. For the KO- IV vs IDS-KO comparison, Prkab2, Uba5, Tacc2, Smpd1, and Scfd1 were upregulated, and Xirp1, Rplp1, Dmac2l, Phka1, and Eef1e1 were downregulated. For the KO-NEB vs IDS-KO, Prkab2, Uba5, Vti1b, Gadd45gip1, and Ssr1 were upregulated, and Bsg, Anxa1, Arhgap1, Mbnl2, and Atp6v1a were downregulated.

### 3.2 Group Variance and Effects

The PCA and MDS analyses were used to evaluate whether alterations in the proteomic profile were consistent within each experimental group and to assess differences in sample organization across groups. PCA separated samples according to the main sources of variation in protein abundance, facilitating visualization of group separation and potential outliers. The first two principal components explained approximately 65% of the total variance (**Figure 3-A**). In this representation, KO-NEB and WT samples were separated mainly along PC1, whereas KO and KO-IV clustered closely together in both principal components. The global heatmap of the 928 significant proteins (**Figure 3-C**) further demonstrated that biological replicates within each experimental group exhibited highly similar abundance profiles, while distinct proteomic signatures were observed across the four groups. To complement the PCA, metric MDS was performed using the pairwise Euclidean distance matrix. Unlike PCA, which maximizes explained variance, MDS seeks to preserve pairwise dissimilarities among samples in a two-dimensional representation. The MDS projection revealed a clear separation between KO-IV and untreated KO samples that was not apparent in the PCA projection (**Figure 3-B**), indicating that treatment-associated differences extend beyond the dominant sources of variance captured by PCA. Consistent with these observations, the pairwise Euclidean distance matrix showed short within-group distances across all experimental groups (**Figure 3- D**), supporting the reproducibility of biological replicates. Moreover, Euclidean distances from the untreated KO baseline were significantly greater for KO-NEB than for KO-IV (47.5 ± 3.7 vs. 40.8 ± 1.1; two-sided Mann-Whitney U test, p < 0.00407), demonstrating that adjunctive nebulization produced a larger global proteomic shift than intravenous treatment alone. PCA, MDS, the global heatmap, and pairwise Euclidean distance analyses consistently demonstrated that the observed proteomic differences reflect robust treatment-associated biological changes rather than within-group variability.

**Figure 3.**
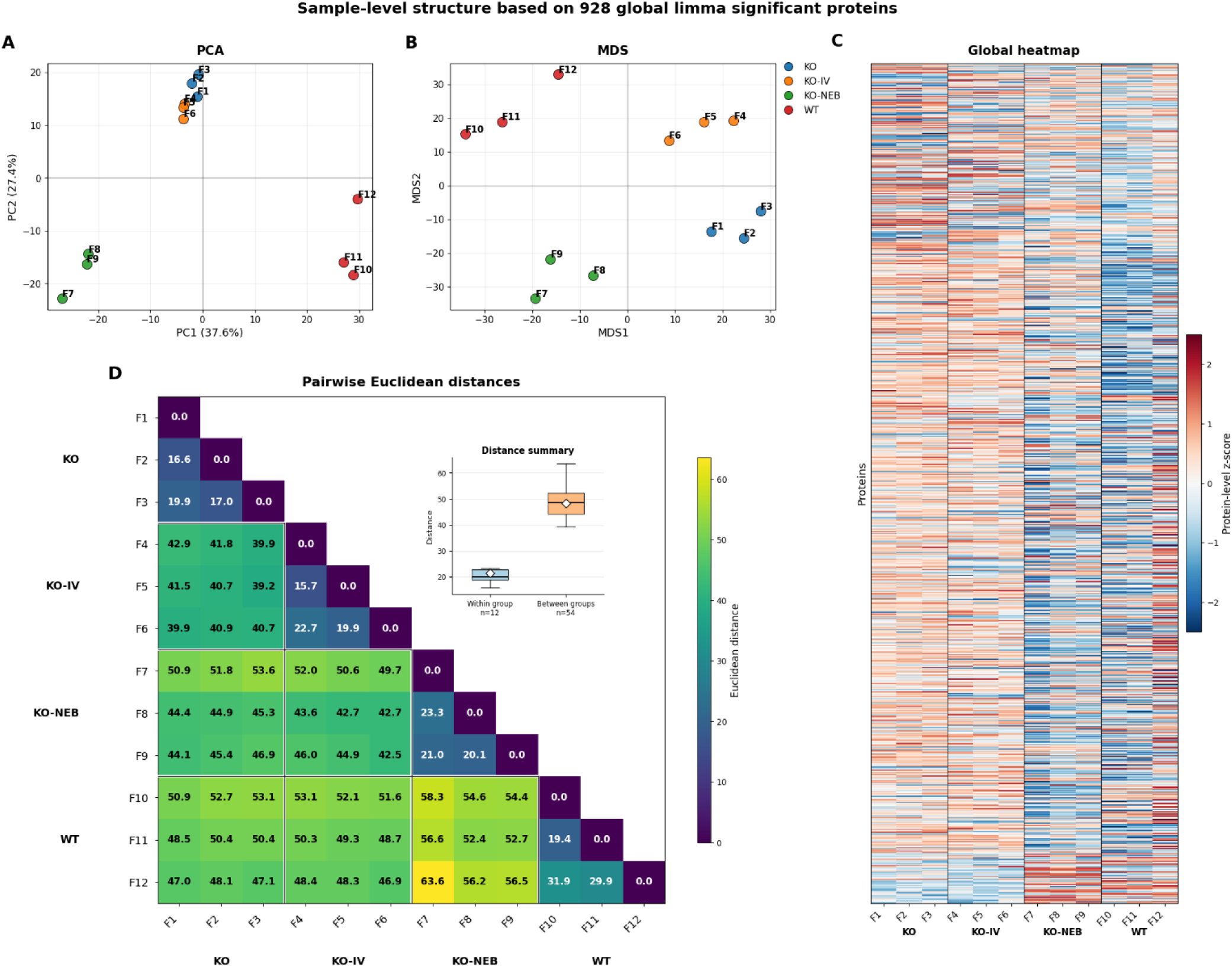
Sample-level structure of the cardiac proteome based on 928 global limma significant proteins. (A) Principal component analysis (PCA) of the 12 cardiac samples using the 928 proteins identified as significant in the global limma moderated F-test. Each point represents one sample, colored according to experimental group. The first two principal components summarize the main sources of proteomic variation among untreated IDS-KO, KO-IV, KO-NEB, and WT samples. (B) Multidimensional scaling (MDS) of the same 12 samples based on pairwise Euclidean distances calculated from the standardized abundance profiles of the 928 proteins. MDS provides a complementary distance-based representation of sample relationships and reveals clearer separation between KO and KO-IV samples than observed in the PCA projection. (C) Global heatmap of the same 928 proteins across all samples. Protein abundance values are shown as protein-level z-scores, with red indicating higher relative abundance and blue indicating lower relative abundance. Vertical lines separate the experimental groups. (D) Pairwise Euclidean distance heatmap calculated from the standardized 928-protein abundance matrix. Only the lower triangle is shown, with colors representing the magnitude of sample-to-sample proteomic distances according to the indicated scale. The inset boxplot summarizes within-group and between-group Euclidean distances, providing an overview of sample similarity within experimental groups relative to distances across groups.

### 3.3 Subcellular Compartment Targets

The outcomes from PCA, MDS, t-SNE, and UMAP analyses by compartment and dominant group highlight the potential cellular- and protein-level signatures of disease and treatments, based on their spatial organization of altered proteins (**Figure 4**). Because individual proteins may contain localization terms from more than one category, we assigned a single main compartment only for visualization purposes using a pre-defined list: Mitochondrion, plasma membrane, endoplasmic reticulum, golgi, endosome, ERGIC/cis-Golgi, lysosome, peroxisome, actin-related, and protein complex. Proteins that did not match any of these categories were labeled as other/unannotated.

**Figure 4.**
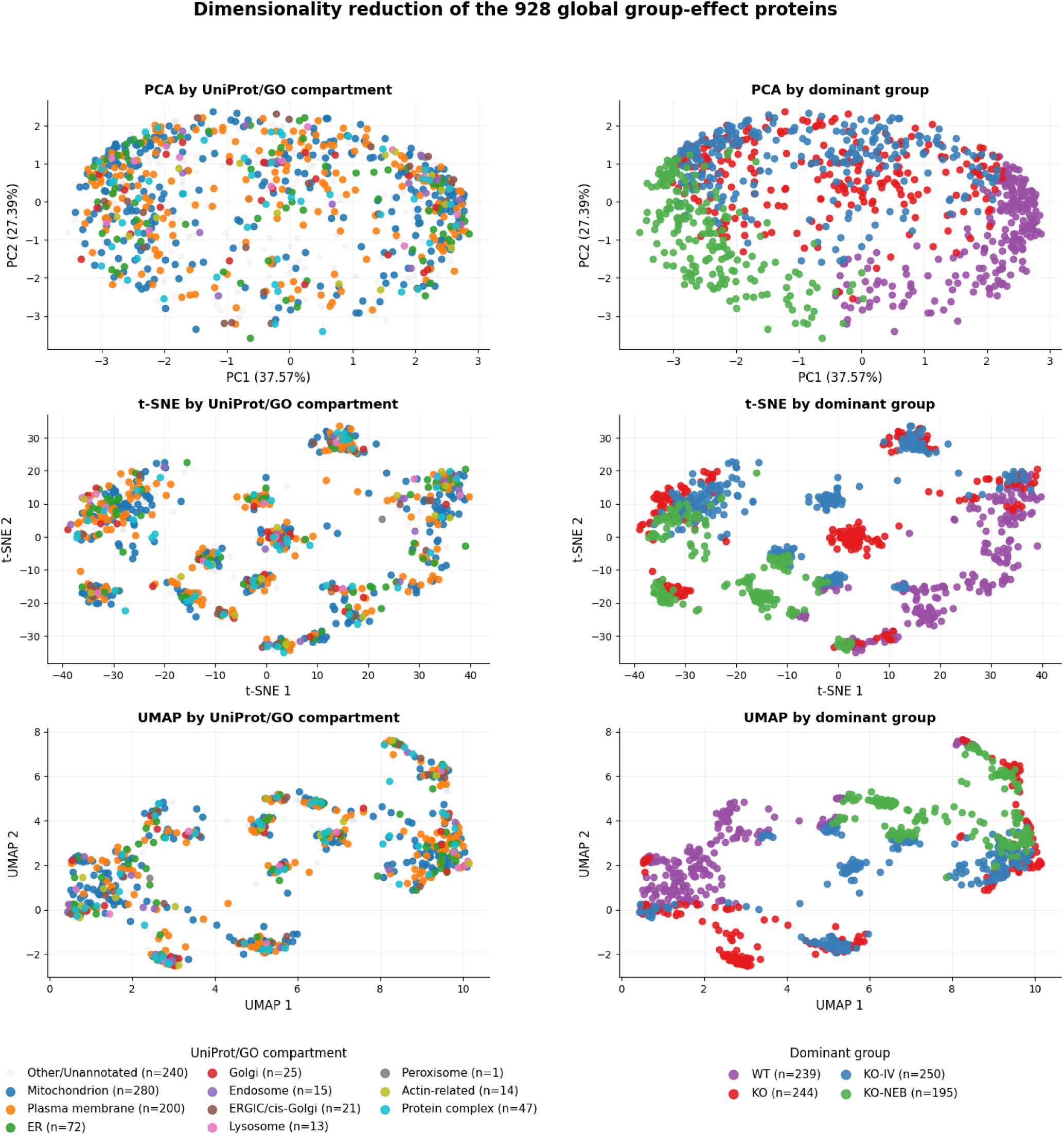
Dimensionality reduction of the 928 global group-effect cardiac proteins accordin to subcellular compartment and dominant experimental group. Principal component analysis (PCA; A and B), t-SNE (C and D), and UMAP projections (E and F) of the 928 proteins identified as significant in the global limma moderated F-test. In the left column, proteins ar colored according to their assigned main UniProt/GO subcellular compartment category, including mitochondrion, plasma membrane, endoplasmic reticulum, Golgi, endosome, ERGIC/cis-Golgi, lysosome, peroxisome, actin-related proteins, protein complexes, and Other/Unannotated proteins. In the right column, proteins are colored according to the experimental group in which they showed the highest mean abundance, defined here as the dominant group. These complementary projections highlight the spatial organization of altered proteins and suggest compartment- and group-associated proteomic patterns across WT, untreated IDS-KO, KO-IV, and KO-NEB cardiac samples.

The highest values across the abundance range of the 928 altered proteins were detected in proteins assigned to the plasma membrane, mitochondrion, actin-related, and protein complex categories (**Figure 5-B**). The radar plot in **Figure 5-A** shows the mean standardized abundance profile of each compartment across groups. The lysosome protein compartment profile of the KO-NEB appeared closer to WT in comparison to the KO and KO- IV groups, which suggests directional change toward normalization within the time frame studied that is highly relevant because IDS-KO is a pre-clinical model for lysosomal storage disorder. Additionally, the compartment group analysis revealed that the overall compartment profile in KO-NEB shifted toward the WT profile, whereas this pattern was less evident in KO- IV.

**Figure 5.**
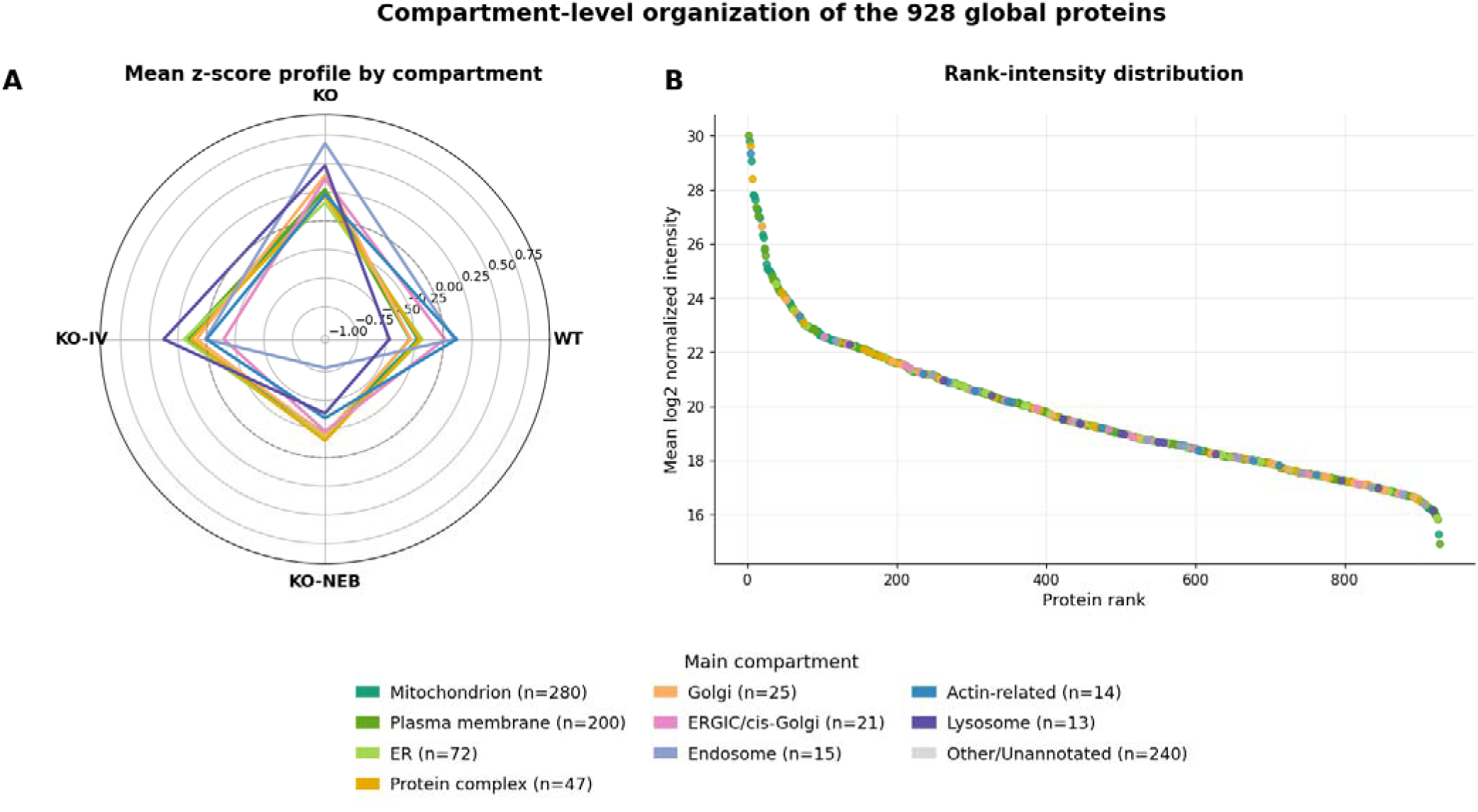
Compartment-level organization and abundance distribution of the 928 global group- effect proteins. (A) Radar plot showing the mean protein-level z-score profile of the 928 global limma significant proteins summarized by main UniProt/GO subcellular compartment across WT, untreated IDS-KO, KO-IV, and KO-NEB groups. Each line represents a compartment category, and the values indicate the mean standardized abundance profile of proteins assigned to that category. Peroxisome was excluded from the panel because it contained only one protein. Rank-intensity plot of the same global protein set. (B) Proteins were ranked by their mean log2-normalized intensity across all samples, with each point representing a protein and colors indicating the assigned main compartment. Other/Unannotated proteins are shown in gray. The distribution indicates that compartment-associated proteins are represented across the abundance range of the altered cardiac proteome, rather than being restricted to highly abundant proteins.=

### 3.4 Protein Clusters

The clustering analysis organized the 928 proteins with significant group-associated variation in abundance per module into 6 groups (clusters, C1-C6). C1 contains 108 proteins and shows low abundance in KO, increased abundance in KO-IV and KO-NEB, and intermediate/low abundance in WT. C2 contains 127 proteins and shows the highest abundance in KO-IV, moderate abundance in KO, and reduced abundance in KO-NEB and WT. C3 contains 134 proteins and shows the highest abundance in KO-NEB, reduced abundance in KO-IV, and intermediate abundance in KO and WT. C4 contains 244 proteins and showed the highest abundance in WT and the lowest abundance in KO-NEB. C5 contains 229 proteins and shows high abundance in KO, KO-IV, KO-NEB, but low abundance in WT. Finally, C6 contains 86 proteins and shows high abundance in KO but reduced abundance in WT, KO-IV, KO-NEB. The top 5 up- and downregulated proteins and their main function in each cluster are described in **Table 1**. Table 1 highlights the functional divergence of the treatment responses, demonstrating that KO-NEB specifically induces proteins involved in mitochondrial network remodeling (e.g., Mfn2 in C1) while suppressing disease-associated pathways. Notably, nebulized ERT successfully reversed the upregulation of proteins that may drive pathological cardiac remodeling and hypertrophy (e.g., Titin and Akt1 in C6), indicating a more comprehensive rescue of the cardiac physiological baseline compared to IV administration alone.

**Table 1.**
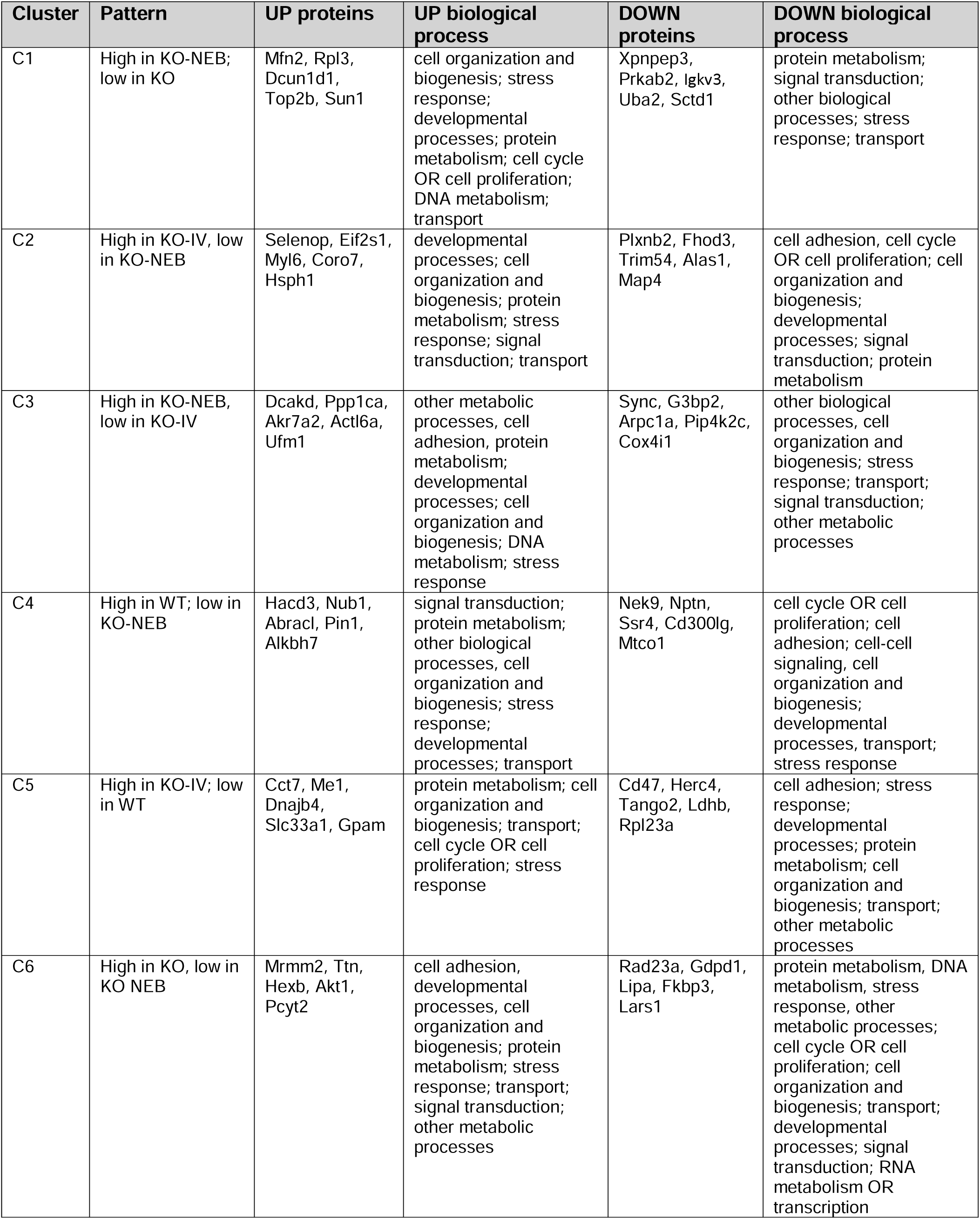
Top 5 up- and down-regulated proteins, main biological processes and patterns in each cluster of proteins.

## 4 Discussion

In the current study, we examined changes in the cardiac proteome caused by IDS-KO and the impact of two different approaches of Idursulfase ERT (intravenous vs. adjunctive nebulization). Our proteomics analyses identified potential protein and subcellular compartment targets contributing to cardiac dysfunction in IDS-KO. The main finding is that idursulfase delivered by nebulization as adjunctive to IV enhances the impact on the cardiac proteome with changes that are consistent with a profile that more closely associated with WT than IV treatment alone.

### 4.1 Protein Candidates and Potential Therapeutic Targets

The abundance analysis suggests potential protein targets contributing to cardiac dysfunction in IDS-KO (**Figure 1).** For example, Ubiquitin-Like Modifier Activating Enzyme 5 (Uba5) is downregulated in IDS-KO but rescued by treatments. Uba5 is involved in the first catalytic step of Ufmylation [24, 25], which plays critical roles in ribosome cycling and DNA damage repair [26, 27]. Additionally, Protein Kinase AMP-Activated (AMPK) Non-Catalytic Subunit Beta 2 (Prkab2) is also downregulated in IDS-KO but rescued by treatments. Prkab2 is involved in AMPK regulation, which senses low ATP levels and activates downstream signaling pathways responsible for stimulating ATP production from protein, lipid and carbohydrates stores [28]. AMPK is crucial for lysosomal metabolism and regulation of mitochondria biogenesis and bioenergetics [28, 29], which are compromised in MPS-II models [30].

Mutations in genes encoding regulatory subunits of AMPK complex, such as Prkag2 and Prkab2, lead to cardiac abnormalities [31–33]. For example, Prkag2 syndrome is a glycogen storage disease characterized by a missense mutation in the Prkag2 gene that can lead to congenital heart failure, atrial fibrillation and cardiac death [32]. Although more robust evidence exists linking Prkag2 deficiency and cardiac dysfunction, a previous clinical report has confirmed the involvement of a deletion in the Prkab2 subunit gene in patients diagnosed with Tetralogy of Fallot—a condition characterized by severe ventricular septal defect, impaired blood flow and cardiac hypertrophy [33, 34].

The deleterious impact of AMPK complex subunit deficiency on the heart and on lysosomes combined with our results showing its downregulation in IDS-KO and its rescue with Idusulfare ERT treatments indicates a potential role of Prkab2 in the cardiac dysfunction phenotype in IDS-KO.

### 4.2 Impact of Idursulfase on Global Cardiac Proteome in IDS-KO

The cardiac proteome changed in a generally similar fashion in both KO-IV and KO- NEB groups. However, adjunctive nebulization (KO-NEB group) nearly doubled the amount of protein abundance changes compared to IV alone, 165 versus 85 proteins **(Figure 2)**. In addition, KO-NEB and WT groups shared more protein abundance changes than KO-IV and WT, 75 versus 59. The overall sample-level organization further supports this observation. PCA revealed a clear separation among the four experimental groups, whereas metric MDS, which preserves pairwise Euclidean distances between samples, provided a complementary representation that highlighted the distinct positioning of the KO-IV and KO-NEB groups relative to untreated KO **(Figure 3A-B**). These multivariate analyses were fully consistent with the pairwise Euclidean distance matrix and the global proteomic heatmap (**Figure 3C-D**), demonstrating that the observed proteomic signatures reflect robust biological differences rather than within-group variability. These findings indicate that adjunctive Idursulfase nebulization enhances the effects of intravenous ERT by producing a larger global remodeling of the cardiac proteome toward the WT state.

### 4.3 Impact of Idursulfase on Subcellular Compartment Profile in IDS-KO

We expanded our protein analyses to identify subcellular compartments affected by IDS-KO and treatments **(Figure 4)**. The KO-NEB group presented a lysosome mean z-score compartment profile very similar to the WT group, while the KO-IV group presented a lysosome mean z-score similar to the IDS-KO group. Both KO-IV and KO presented distinct lysosome compartment profiles compared to KO-NEB and WT (**Figure 5**). These findings are relevant because MPS-II is a lysosomal storage disease and our data suggests that nebulized Idursulfase ERT adjunctive to intravenous delivery optimized the shift in the protein profile of cardiac lysosomes of IDS-KO mice towards that seen in WT.

The compartment group analysis also revealed that endosome protein compartment changes caused by IDS-KO were restored in the KO-NEB but not in the KO-IV treatment group (**Figure 5**). Endosomes are responsible for transporting cellular waste to lysosomes, where it is degraded. In lysosomal storage disorders, impairments in lysosomal function can stimulate endosome activity and proteomic changes due to the accumulation of cellular waste [35]. The similar lysosome proteome profile between WT and KO-NEB and the opposite endosome proteome profile between KO and KO-NEB suggests that adjunctive nebulized Idursulfase ERT modulates the proteome signature of cellular compartments in the heart that are heavily affected by IDS-KO.

### 4.4 Modulation of Protein Clusters by IDS-KO and Idursulfase ERT

The purpose of the novel clustering analysis was to identify groups of proteins with a similar abundance pattern in specific experimental groups. The clustering analysis identified coordinated proteomic modules associated with disease status and treatment responses. The cluster-compartment association analyses of altered proteins demonstrate that all 6 clusters exhibit similar subcellular compartment signatures: higher proportion of mitochondrial proteins, followed by plasma membrane, endoplasmatic reticulum (ER) and protein complex. (**Figure 7**).

**Figure 6.**
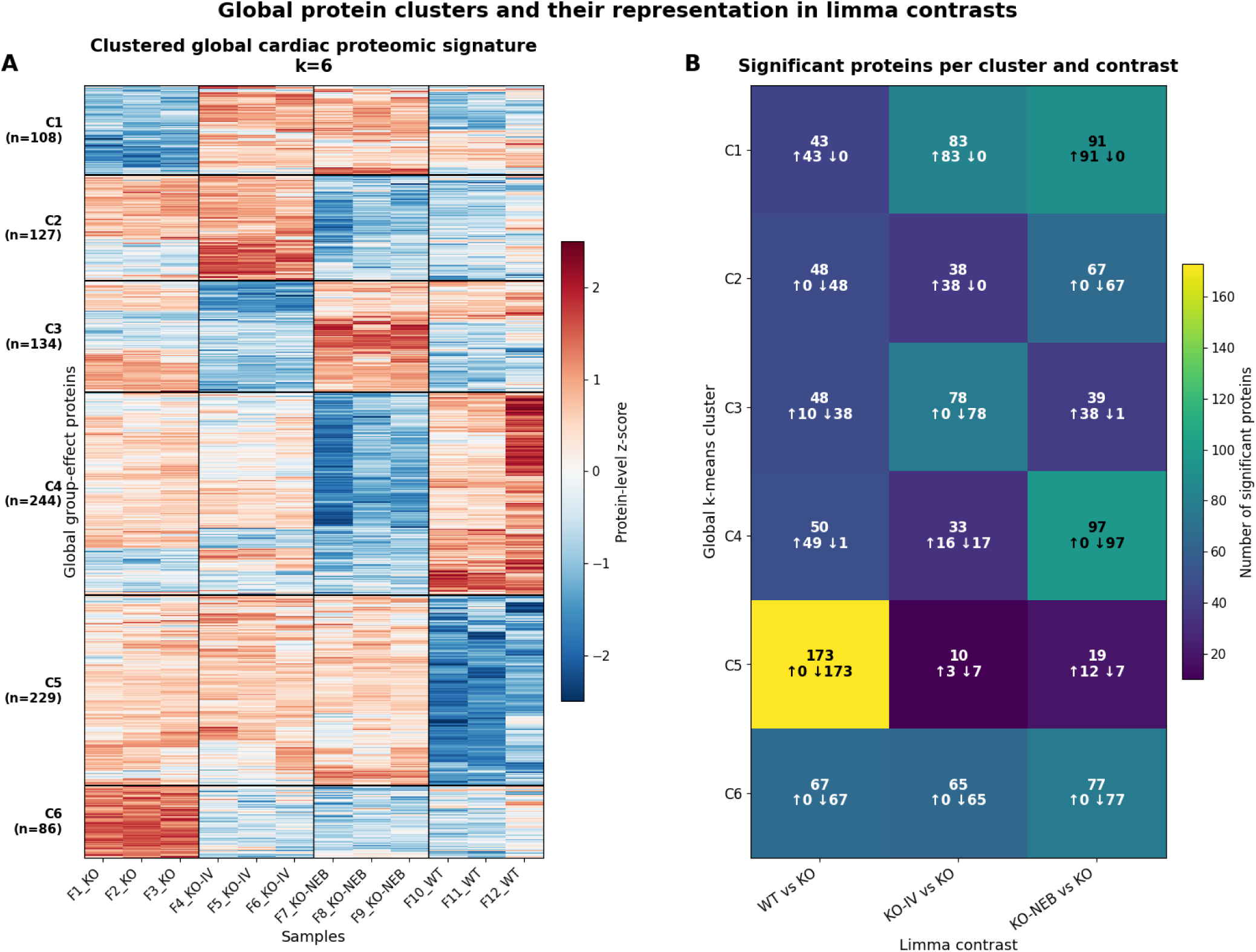
Global cardiac proteomic modules and their contribution to treatment- and disease- associated contrasts. (A) The left heatmap shows k-means clustering of the 928 global group- effect proteins identified by limma moderated F-test. Rows represent proteins and columns represent individual cardiac samples. Values are protein-level z-scores calculated across samples, with red indicating higher relative abundance and blue indicating lower relative abundance. Proteins were grouped into six clusters (C1–C6), representing coordinated abundance modules across KO, KO-IV, KO-NEB, and WT samples. (B) The right heatmap shows the number of significantly altered proteins from each cluster represented in the planned pairwise limma contrasts. Each cell reports the total number of significant proteins, followed by the number of upregulated and downregulated proteins. Differential abundance was defined as |log2FC| ≥ 1 and adjusted p-value < 0.05.

**Figure 7.**
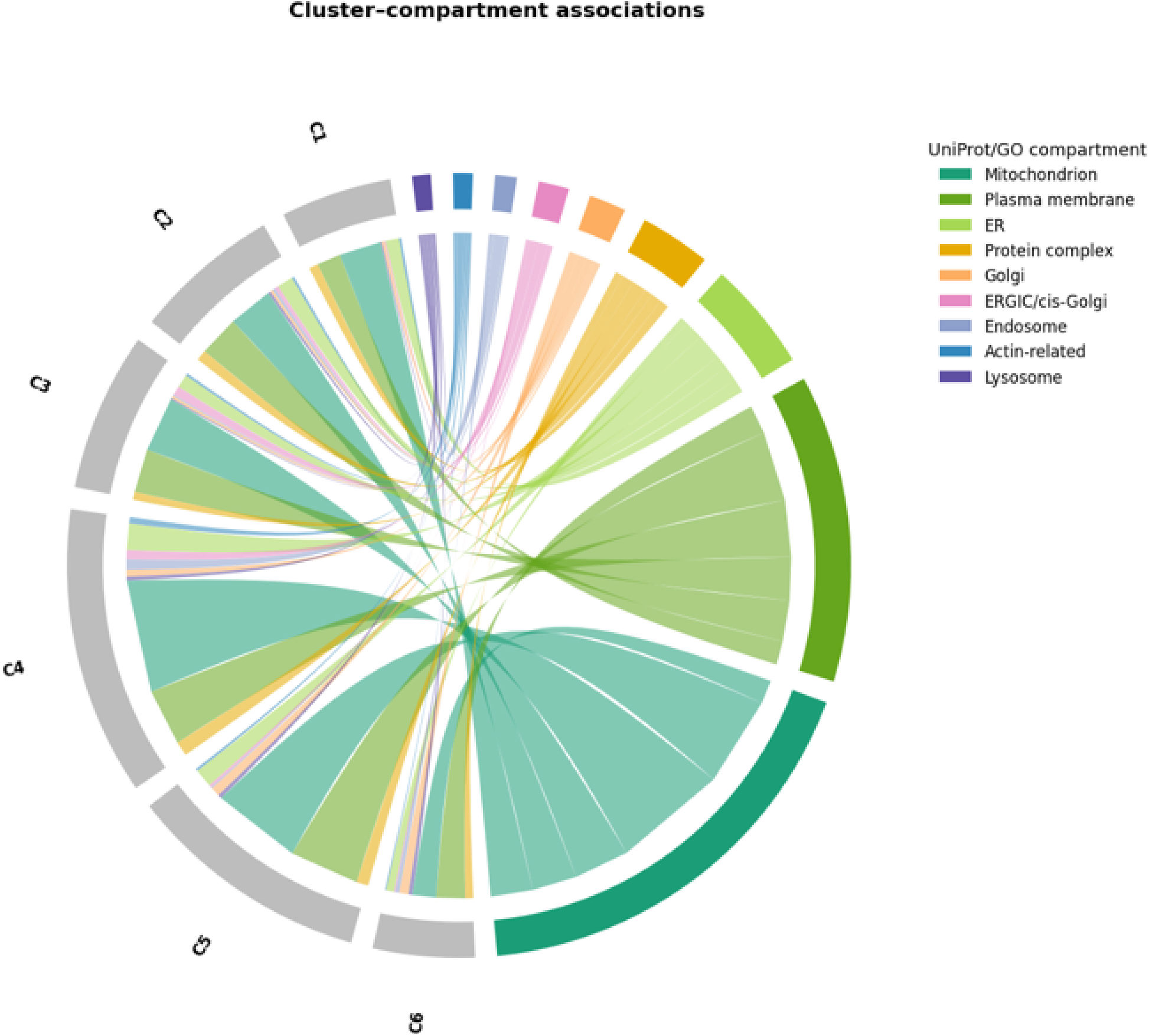
Cluster–compartment associations among global cardiac proteomic modules. Chord diagram showing the relationship between the six k-means protein clusters (C1–C6) and the main UniProt/GO subcellular compartment categories assigned to the 928 global group-effect proteins. Gray sectors represent protein clusters, and colored sectors represent annotated compartments. Ribbon width is proportional to the number of proteins shared between each cluster and compartment. The plot highlights the broad contribution of mitochondrial and plasma membrane proteins across the global cluster structure, with additional representation of endoplasmic reticulum, protein complex, Golgi, ERGIC/cis-Golgi, endosome, actin-related, and lysosome-associated proteins. Other/Unannotated proteins and peroxisome were not shown to facilitate visualization.

All untreated IDS-KO samples presented downregulation and upregulation of proteins clustered in C1 and C6, respectively **(Figure 6)**. Both treatment groups, KO-IV and KO-NEB reversed the overall profile of C1 and C6 proteins. Although C1 reflects proteins induced by both treatment groups, C6 is the most therapeutically relevant module because it reflects proteins elevated in the untreated KO group that are decreased by treatment back to WT levels. Additionally, C2 proteins were preferentially upregulated in KO-IV, but KO-NEB showed a C2 protein profile closer to the WT.

Notably, Titin is one of the most upregulated proteins in C6. Titin controls myocardial stiffness [36], diastolic function [37], and contraction [38], which are all influenced by MPS-II [7]. Titin gene truncations disrupt protein function resulting in dilated cardiomyopathy and heart failure [39]. Additionally, an altered expression ratio of titin isoforms has been reported in dilated cardiomyopathy patients [40–43]; these patients express increased amount of the compliant isoform N2BA. This shift in titin isoform expression is believed to be compensatory and beneficial to counteract cardiomyocyte stiffness and symptoms like exercise intolerance [41]. The increase in titin protein abundance in IDS-KO and the rescue of titin protein abundance profile by Idursulfase ERT may suggest a contributing or a compensatory role of titin in the cardiac adaptations caused by IDS-KO that is worth investigating in future studies.

Together, these results further support distinct proteomic effects of standalone IV or adjunctive nebulization of Idursulfase ERT on the IDS-KO heart and suggest potential targets that can be investigated in future studies.

### 4.5 Adjunctive Nebulization versus standalone IV Drug Administration Considerations

Nebulization is a viable approach for drug delivery to the pulmonary circulation and cardiac muscle [44]. The greater impact of adjunctive nebulization of Idursulfase ERT on the cardiac proteomic signature of IDS-KO mice may reflect a higher dose per animal as well as enhanced absorption and cardiac delivery of Idursulfase through the lungs and pulmonary circulation. Drugs administered via nebulization are rapidly converted in fine particles that are delivered directly to the alveoli; the result is a prompt increase in local drug concentration that is transferred to heart and subsequently pumped to the circulation [18]. Therefore, the pharmacological advantages of nebulized drug administration and the greater dosage used in adjunctive nebulization treatment may explain the greater rescue of the cardiac proteomic signature in IDS-KO by adjunctive nebulized Idursulfase ERT, compared to standalone IV treatment.

## 5 Conclusion

Cardiac dysfunction is a major driver of patient mortality in MPS-II. The current study defined the impact of IDS-KO on the cardiac proteome and whether Idursulfase ERT rescues the cardiac protein changes elicited by IDS-KO, a pre-clinical model of MPS-II. Overall, our results indicate potential protein (e.g. Uba5, Prkab2, and titin) and subcellular compartment (e.g. lysosomes, endosomes, and mitochondria) targets involved in the cardiac pathophysiology with deficiency of IDS. The main finding is that adjunctive nebulized Idursulfase ERT is more effective than IV alone in shifting the cardiac proteome signature of IDS-KO mice toward WT.

## 6 Acknowledgments

The study was funded by an investigator-initiated grant from Takeda Pharmaceuticals (IIR-USA-001523). We thank Dr. Kari Basso from the University of Florida for her support throughout the project. The University of Florida Proteomics and Mass Spectrometry Core was funded by NIH grants S10 OD021758-01A1 and S10 OD030250-01A1 that partially supported execution of this study. The authors also sincerely thank William Skelly for suggesting the limma package and multi-dimensionality scaling for analysis of our data.

